# The table-top visual search ability test for children and young people: normative response time data from typically developing children

**DOI:** 10.1101/824672

**Authors:** Jonathan Waddington, Jade Pickering, Timothy Hodgson

## Abstract

Five table-top tasks were developed to test the visual search ability of children and young people in a real-world context, and to assess the transfer of training related improvements in visual search on computerised tasks to real-world activities. Each task involved searching for a set of target objects among distracting objects on a table-top. Performance on the Table-top Visual Search Ability Test for Children (TVSAT-C) was measured as the time spent searching for targets divided by the number of targets found. 108 typically developing children (3-11 years old) and 8 children with vision impairment (7-12 years old) participated in the study. A significant correlation was found between log-transformed age and log-transformed performance (*R*^2^ = 0.65, *p* = 4 × 10^−26^) in our normative sample, indicating a monomial power law relationship between age and performance with an exponent of −1.67, 95% CI [−1.90, −1.43]. We calculated age-dependent percentiles and receiver operating characteristic curve analysis indicated the 3^rd^ percentile as the optimal cut-off for detecting a visual search deficit, giving a specificity of 97.2%, 95% CI [92.2%, 99.1%] and sensitivity of 87.5%, 95% CI [52.9%, 97.8%] for the test. Further studies are required to calculate measures of reliability and external validity, to confirm sensitivity for visual search deficits, and to investigate the most appropriate response modes for participants with conditions that affect manual dexterity. Additionally, more work is needed to assess construct validity where semantic knowledge is required that younger children may not have experience with. We have made the protocol and age-dependent normative data available for those interested in using the test in research or practice, and to illustrate the smooth developmental trajectory of visual search ability during childhood.

## Introduction

Search is a fundamental human skill involved in performing a wide range of daily tasks and achieving operational goals, such as finding a favourite book on a bookshelf lined with other books. Searching for an object defined by one or more perceptual features (e.g. colour, size, or shape) requires attentional processes to actively scan the environment for the features that define the target object relative to distracting objects. Computerised visual search tasks have been used to study this ability in healthy adults and children but many aspects of the development of visual attention in children remain poorly understood (Johnson, 2019). Studies that have examined visual search ability across the lifespan have revealed an inverted U-shaped performance pattern with performance increasing from childhood into adulthood and declining from middle age (Plude, Enns & Brodeur, 1994). Increased efficiency in processes that integrate the perceptual features of objects and orient the focus of attention between locations and objects are thought to drive improvements in search ability between childhood and adulthood; whereas decreased efficiency in attentional orienting alone is thought to be the cause of decline in later life (Trick & Enns, 1998).

The visual search ability of children depends on the presence or absence of oculomotor impairments, visual field impairments and attentional impairments such as an increased influence of multiple distracting objects in close proximity or “crowding” (Huurneman, Cox, Vlaskamp & Boonstra, 2014). Children’s ability to organise complex visual search strategies efficiently can also be limited by the development of executive functions such as working memory, interference control, and planning (Woods et al., 2013). Some adults with visual field impairments (i.e. hemianopia) present with inefficient search strategies as indexed by eye movement recordings, whilst others develop compensatory search strategies that enable them to perform visual search almost as effectively as adults without visual field impairments (Zihl, 1995). The factors that determine development of compensatory search strategies remain unclear, and cannot be explained simply as a matter of time since or extent of the visual field deficit (Zihl, 1999, 2000). However, the observation that compensatory search strategies can be developed spontaneously has led to the development of visual search training and rehabilitation programmes designed to promote compensatory search strategies in practice (e.g. Turton et al., 2015).

Previous research into visual search training has focused on improving visual search skills in adults with either hemianopia or spatial inattention caused by stroke. Many modern training protocols designed for this purpose involve actively searching for digital targets displayed on a screen (e.g. Roth et al., 2009; Aimola et al., 2014; Ong et al., 2015; Sahraie, Smania & Zihl, 2016; Chesham et al., 2019). Current research projects are also focusing on developing training protocols to improve visual search skills in children with visual field impairments as a consequence of perinatal and acquired brain injury (Waddington & Hodgson, 2017; Ivanov et al., 2018; Waddington et al., 2018), or as a consequence of surgical interventions to treat other conditions such as hemi-disconnection surgery to treat intractable epilepsy (Koenraads et al., 2014).

Previous studies have demonstrated that training-related improvements in visual search are specific and task dependent (Schuett, Heywood, Kentridge, Dauner & Zihl, 2012). Training skills using a digital platform may only deliver context specific improvements which are confined to the task and medium within which the training took place (e.g. digital vs. real-world), particularly for younger children (Huber et al., 2016; Zack & Barr, 2016). Whilst there are many benefits to using a digital format to assess visual cognitive abilities (e.g. improved measurement accuracy, portability with mobile platforms, and greater control of image presentation), there is a need to evaluate generalisation of training-related improvements in visual search from a digital format to real-world scenarios. Such a test would be able to give objective outcome measures of visual search ability without actively searching for digital targets displayed on a screen. Table-top assessment tools are established in neuropsychological testing for vision impairment (e.g. Brown, 2016; Brown & Peres, 2018a, 2018b; Cooke, McKenna, Fleming & Darnell, 2006; Donnelly, Hextell & Matthey, 1998; Katz, Itzkovich, Averbuch & Elazar, 1989; Reitan & Wolfson, 2004; van Vugt, Fransen, Creten & Paquier, 2000). However, the majority of these tests are based on probing specific visual and cognitive functions and are not designed to assess functional vision or meaningful vision-related activities with real-world objects, and many have not been validated for use with children.

We have developed a battery of five table-top tasks to measure visual search ability in children. The Table-top Visual Search Ability Test for Children, or the TVSAT-C, was designed to be suitable for use by professionals at relatively low cost with readily available and portable household objects. It was also designed to be used in conjunction with other tests, observations, and history taking as part of a comprehensive functional vision assessment rather than as a single vision screening measure or to test a specific visual function. The results of a comprehensive functional vision assessment can be used to individualise support plans or interventions for children and young people with vision impairment, in both education and habilitation contexts. The TVSAT-C has already been piloted to evaluate changes in the visual search ability of children and young people with homonymous visual field impairments before and after a six-week visual search training programme (Waddington et al., 2018). The aim of the present study was to investigate how age affected visual search performance on the TVSAT-C with a normative sample of children. A secondary aim was to investigate whether these reference values could be used to detect deficits in visual search ability with a pilot sample of eight vision impaired children with a range of medical conditions.

## Methods

### Participants

This study received ethical approval from the University of Lincoln’s School of Psychology Research Ethics Committee. We identified candidates for participation during a public engagement event held at the University of Lincoln. Children from 3 to 11 years old and their parents were invited to attend the five-day event for one morning or afternoon, to take part in a variety of research studies designed to be engaging for children and young people. In this particular study we included any child that expressed an interest in participating and excluded the data for children who self-reported a vision impairment or other special educational needs. 214 candidates attended the event and 122 participants assented with parental consent. We excluded the data from 14 participants who reported special educational needs and analysed the data from 108 participants (55 female; median age = 6.4 years, interquartile range = 5.1 – 7.7 years, absolute range = 3.1 – 10.5 years).

Comparative data were also obtained from 8 participants with diagnosed vision impairment (7-12 years old). These participants were identified from specialist schools, colleges, and units during two separate studies to develop and evaluate a therapeutic game, designed to promote visual search training for young people with homonymous visual field impairment (Waddington, Linehan, Gerling, Hicks & Hodgson, 2015; Waddington et al., 2018). Those studies received ethical approval from the University of Lincoln’s School of Psychology Research Ethics Committee and the United Kingdom National Research Ethics Service Committee North East – Newcastle & North Tyneside 1. Five participants were diagnosed with visual field impairment due to brain injury (hemianopia or quadrantanopia confirmed by perimetry), two were diagnosed with albinism, and one was diagnosed with congenital fibrosis of the extraocular muscles. Participants with visual field impairment did go on to complete visual search training but we present their initial test results, prior to training, here.

### Tasks, Materials and Protocol

The TVSAT-C tasks were adapted from a test previously used to evaluate outcome measures after visual search training in adults with hemianopia (Pambakian, Mannan, Hodgson & Kennard, 2004). The original test was designed to assess functional vision (i.e. how well the person functions during vision-related activities) as opposed to assess a specific visual function (i.e. how well a component of the visual cognitive system functions). As such, the tasks involved real-word activities that required some manual dexterity and intelligence to perform. Changes from the previous test were designed to make the tasks more age-appropriate and functional for children. For example, we adapted one task that involved threading beads bearing letters of the alphabet onto a leather lace in alphabetical order to a task that involved correctly matching brightly coloured magnets in the shape of letters of the alphabet to a series of letters written on a card. The tasks ranged in difficulty such that more difficult tasks in the assessment battery would challenge the more able participants and improve sensitivity of the test at that end of the spectrum of ability, while less able participants would still be able to complete the less difficult tasks. We also selected test materials that were reasonably large, bold and plain-coloured, as well as relatively easy to pick up and manipulate. Five timed search tasks were included in the test to keep the assessment period short in duration and maintain engagement.

The materials required to perform the TVSAT-C included one Attribute Blocks Desk Set (Learning Resources; Illinois, USA; LER 1270; https://www.learningresources.com/attribute-blocks-desk-set) that originally contained 60 plastic blocks: a combination of five shapes, two sizes, two thicknesses, and three colours. The hexagon shapes and small sized blocks were excluded, leaving 24 attribute blocks in the test kit. The test kit also required one set of Wooden Letter Alphabet Magnets (Melissa & Doug; Connecticut, USA; MAD1111255E1; https://www.melissaanddoug.com/wooden-letter-alphabet-magnets/448.html) that originally contained 52 upper- and lowercase magnetic letters in blue, green, red and yellow. Only the uppercase letters were included in the test kit. We also used a modular Compact Disc (CD) Storage Rack with the capacity to hold 120 CDs (Westpoint Design; Castle Douglas, UK; CD120; https://cdanddvdstorage.com/products), eight standard jewel CD cases, and card in four pastel shades (blue, green, red, and yellow). The coloured card was fixed to the clear plastic exterior of the CD cases to create four pairs of coloured CD cases.

The rest of the test kit included 24 UK denomination coins (eight 1p, eight 2p, and eight 5p), one plain black tablecloth, A3 sized (21.0 × 29.7cm) white card, and a black bullet tip marker. Time to complete the tasks was measured using a stopwatch on a smartphone with the assessor manually starting and stopping the stopwatch.

Each task involved searching for one set of target objects among sets of distracting objects on a table covered with a black tablecloth. Task 1 included 3 sets of 8 coloured blocks (blue, red, and yellow) and task 2 included 4 sets of 6 geometric blocks (circles, rectangles, squares, and triangles). Both task 1 and task 2 included the same attribute blocks so there was the same combination of colours and shapes in both tasks (i.e. two blue circles, two blue rectangles, … two yellow squares, two yellow triangles). Task 3 included 3 sets of 8 UK denomination coins (1p, 2p, and 5p). Task 4 included 5 pseudorandomly selected target letters from the modern English alphabet, with the remaining 21 letters acting as distracting objects. Task 5 included 4 sets of 2 coloured compact disk cases (blue, green, red, and yellow). We selected task order and target sets pseudorandomly before each assessment using a simple random number generator (see Table 1 for example).

**Table 1.** Representative test protocol for the TVSAT-C after randomisation of task order and targets.

| Task | All sets (number of objects in each set) | Target set |
| --- | --- | --- |
| 1. Blocks (colour) | Blue (8), red (8), yellow (8) | Blue |
| 2. Blocks (shape) | Squares (6), triangles (6), circles (6), rectangles (6) | Squares |
| 3. Coins | 1p (8), 2p (8), 5p (8) | 5p |
| 4. Letters | Uppercase letters (26) | C T V W I |
| 5. CDs | Blue (2), green (2), red (2), yellow (2) | Red |

We prepared tasks 1-4 by selecting a corresponding diagram of the required targets illustrated on white A3 card. The white A3 card was placed in front of the participant by the assessor with the diagram face down. The sets of target and distracting objects were then placed at random around the perimeter of the white card. At the start of each task the card was turned over to reveal the diagram of the target objects, the assessor then gave a verbal prompt (e.g. “Can you find all the blue objects?” or “Can you find the letters: C, T, V, W, and I?”) to begin the task (see Figure 1 for examples). We prepared task 5 by placing a CD rack (80cm in length) in front of the participant and four CDs, each separated by two compartments on the storage rack, at either end of the rack. The assessor gave a verbal prompt (e.g. “Can you find the red CDs?”) to begin the task. Participants indicated they had found a target by picking it up and placing it in front of them. Each task ended when the participant had placed the last target in front of them or when they indicated that they couldn’t find any more of the targets. The time taken to complete each task and the number of targets found were recorded.

**Figure 1.**
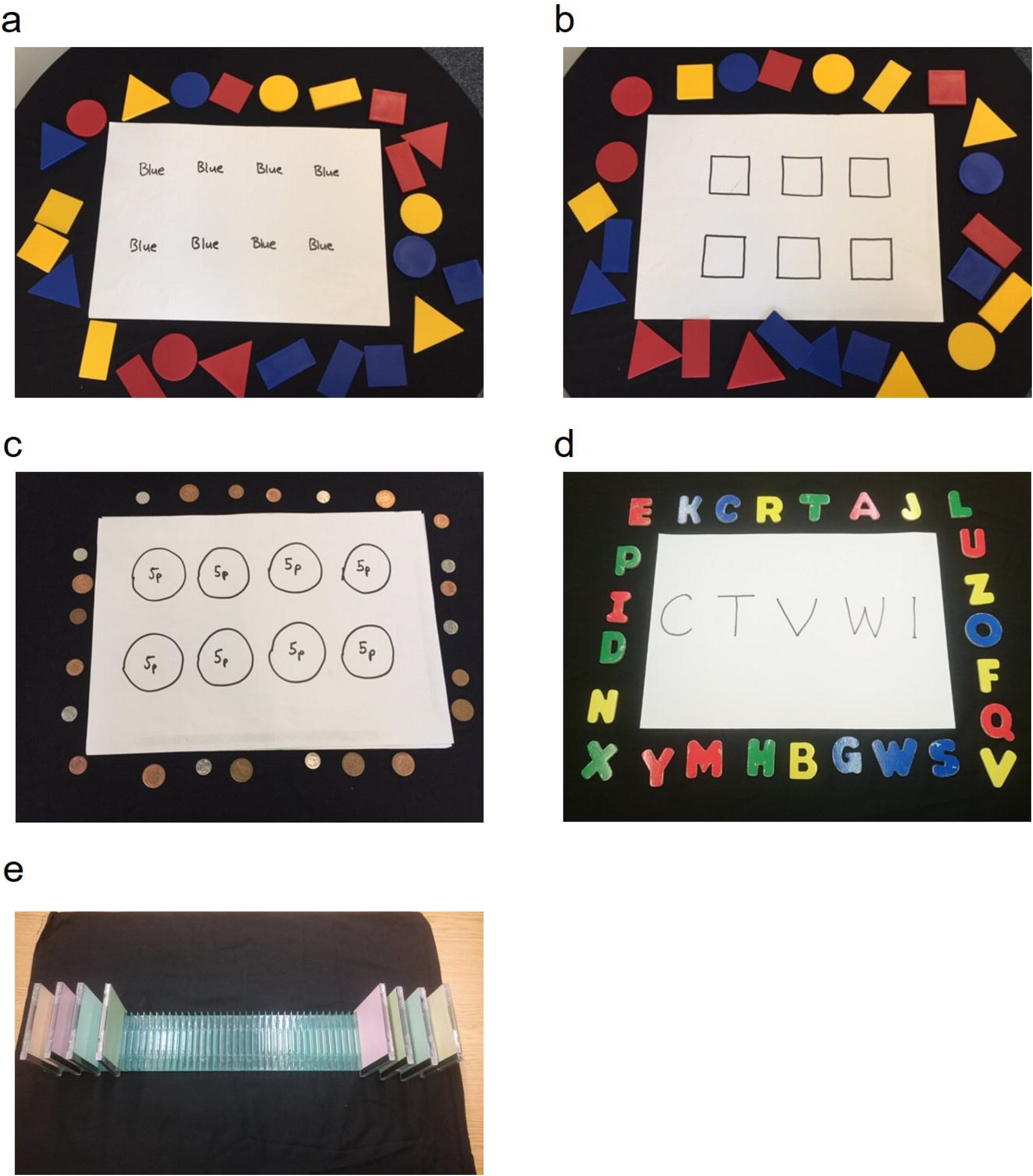
Representative photographs of tasks from the TVSAT-C. Photographs of five representative table-top search tasks from the TVSAT-C, including: (a) 24 attribute blocks with an equal mix of three colours (blue, red, and yellow), (b) 24 attribute blocks with an equal mix of four shapes (squares, circles, rectangles, and triangles), (c) 24 UK denomination coins with an equal mix of three values (1p, 2p, and 5p), (d) 26 letters of the alphabet, and (e) 8 coloured compact disk cases with an equal mix of four colours (blue, green, red, and yellow) presented on a display rack. Note that target objects for the search tasks (a), (b), (c), and (d) are displayed on an A3 sized (21.0 × 29.7cm) white card at the start of the task and represent the tasks displayed in the protocol from Table 1.

### Analysis

We defined the response time performance on the visual search tasks as the total time taken to complete each task in seconds divided by the total number of targets correctly found. In the case where a participant was unable to complete an individual task and no targets were found, no response time was recorded for that task. The distribution of visual response times was significantly different to Gaussian as expected, so visual response time data were logarithmically transformed to approximately conform to normality. We evaluated a simple linear regression model including log-transformed visual response times and log-transformed age. We assessed the residuals of the regression for departure from normality with the Anderson-Darling test. We performed receiver operating characteristic curve analysis to determine the optimal cut-off for detecting potential visual search deficits, and calculated the specificity and sensitivity of the test using the results from our normative and comparative group participants respectively. Confidence intervals for specificity and sensitivity were produced with the Wilson score method (Newcombe, 1998).

## Results

We plotted participants’ visual response times on individual tasks from the TVSAT-C against their age, which indicated similar nonlinear relationships across tasks (Figure 2). We calculated a multivariate linear regression to predict the log-transformed visual response times for each and every task based on log-transformed age. Significant regression equations were found for each and every task with F values ranging from *F*(1,87) = 110 for the Blocks (colour) task to *F*(1,87) = 47.5 for the CDs task (all *p* < 0.001) and adjusted R-squared scores ranging from *R*^2^ = 0.553 to *R*^2^ = 0.346. The coefficients of regression appeared similar across tasks, ranging from −2.03, 95% CI [−2.44, −1.61] for the Letters task to −1.61, 95% CI [−2.08, −1.15] for the CDs task. The log-transformed visual response times from each individual task were also significantly correlated with each other (all *p* < 0.001), ranging from *r* = 0.734 for the Blocks (shape) and Blocks (colour) tasks to *r* = 0.386 for the Coins and CDs tasks. Given this good internal consistency (*α* = 0.869), we combined task results and continued to analyse visual response time performance globally on the TVSAT-C as the sum of time taken to complete every task divided by the sum total of targets correctly found across all tasks.

**Figure 2.**
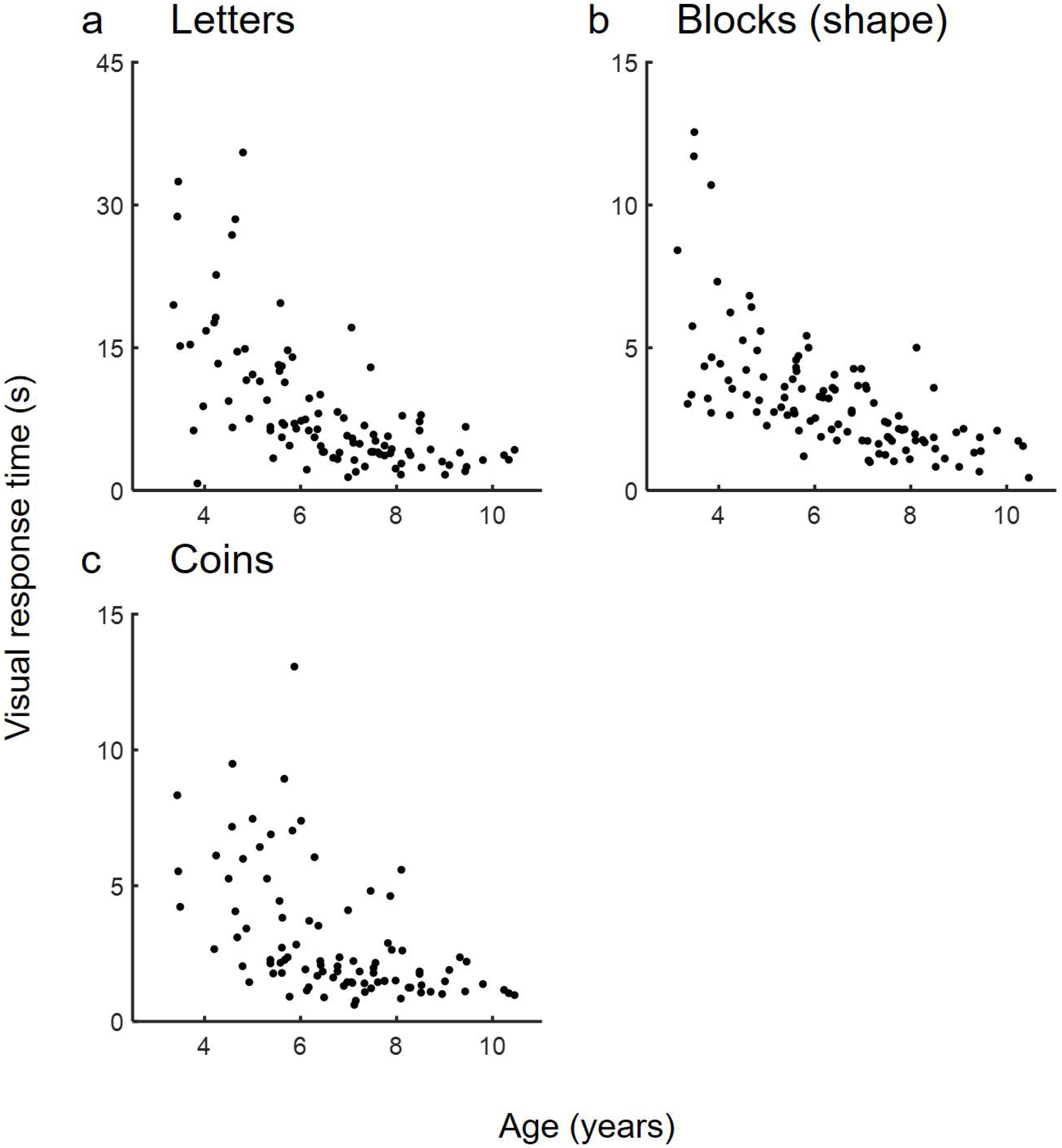
Age-related performance on individual tasks from the TVSAT-C. Plots of performance on individual tasks from the TVSAT-C against age, demonstrating raw data from a normative sample of: (a) 102 children completing the Letters task, (b) 107 children completing the Blocks (shape) task, and (c) 92 children completing the Coins task.

We plotted participants’ global visual response times on the TVSAT-C against their age (Figure 3a), which indicated a nonlinear relationship as expected. The results of Spearman’s rank correlation indicated a significant decreasing monotonic dependency between age and visual response times (*r*_*s*_ (106) = −0.816, *p* = 6.05 × 10^−27^). We calculated a simple linear regression to predict log-transformed visual response times based on log-transformed age. A significant regression equation was found (*F*(1,106) = 200, *p* = 4.00 × 10^−26^) with an adjusted R-squared of *R*^2^ = 0.650, indicating a monomial power law relationship between age and visual response times. After inverting logs, we found that participants’ mean visual response times were equal to exp (4.21) · (*age*)^−1.67^ seconds when age was measured in years. In plain English, participants’ visual response times decreased 16.7% for each 10.0% increase in age, on average. We calculated the confidence interval for the exponent to determine whether the relationship could be approximated by a simple reciprocal relationship and found the exponent to be −1.67, 95% CI [−1.90, −1.43], indicating it could not.

**Figure 3.**
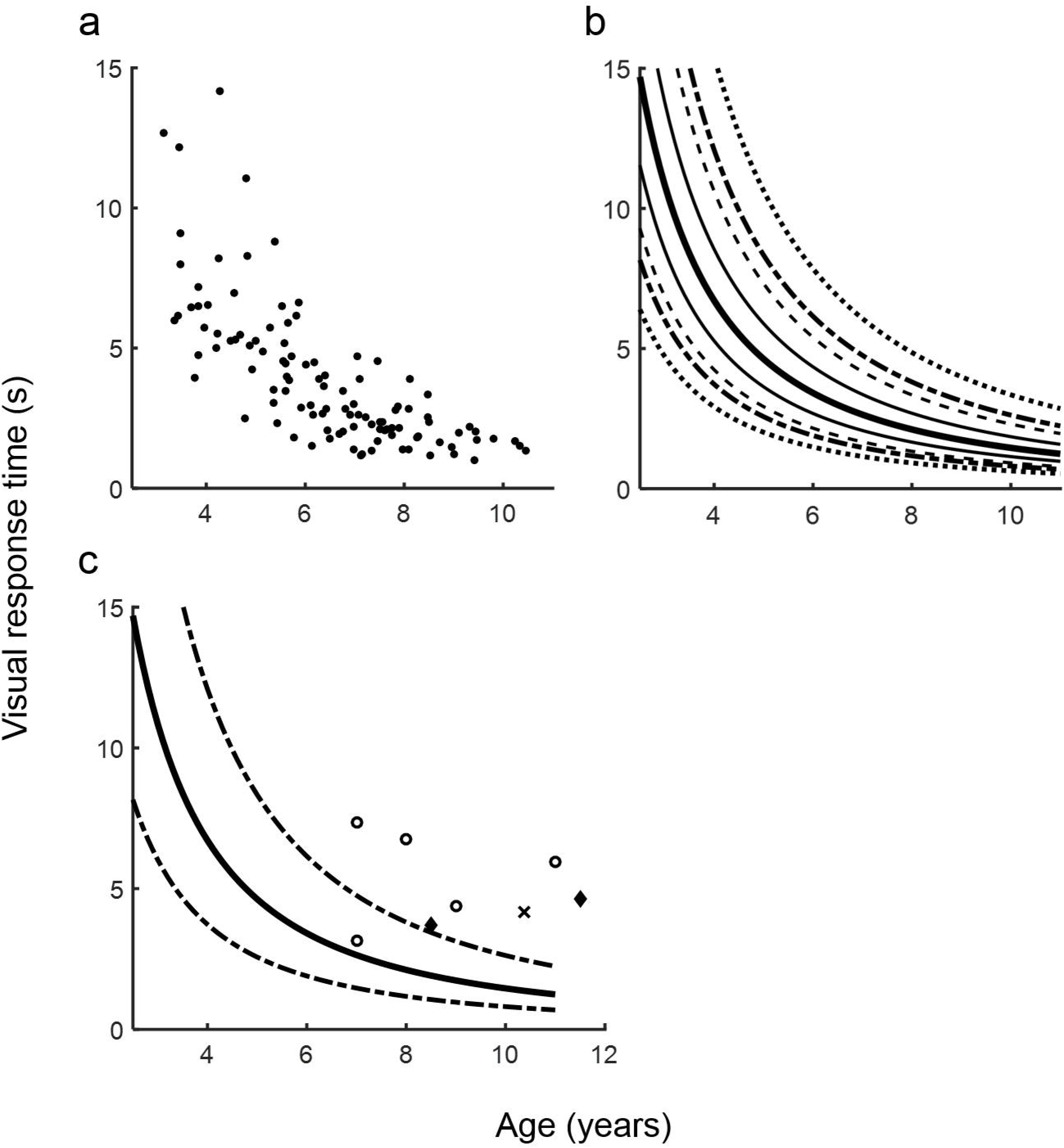
Age-related performance on the TVSAT-C. Plots of performance on the TVSAT-C against age, demonstrating: (a) raw data from a normative sample of 108 children, (b) population percentiles estimated from the normative sample (bold solid line = 50th percentile, thin solid lines = 25th and 75th percentiles, dash lines = 10th and 90th percentiles, dot-dash lines = 5th and 95th percentiles, dot lines = 1st and 99th percentiles) and (c) raw data from a sample of vision impaired children (empty circles = diagnosed visual field impairment due to brain injury, diamonds = diagnosed albinism, cross = diagnosed congenital fibrosis of the extraocular muscles).

We compared the studentised residuals of the linear regression with the predicted values of visual response times and found a symmetrical random distribution, indicating that the assumptions of linearity, additivity, and homoscedasticity had not been violated. An Anderson-Darling test for normality on the standardised residuals found no significant difference between the distribution of standardised residuals and the Gaussian distribution (*A*^2^ = 0.398, *p* = 0.365). We therefore modelled visual response time percentile trajectories with the function:

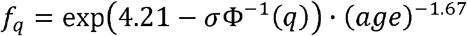

where Φ^−1^(·) is the standard probit function, *σ* is the age-controlled population standard deviation of log-transformed visual response times, and *q* is the desired quantile expressed as a decimal. We estimated *σ* = 0.358 log units from the error variance calculated during the simple linear regression. We plotted the trajectories of the 1^st^, 5^th^, 10^th^, 25^th^, 50^th^, 75^th^, 90^th^, 95^th^, and 99^th^ percentiles for reference (Figure 3b). We calculated the standard z-scores of participants using the model:

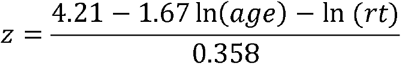

where *rt* is the response time performance measured in seconds per object found, and age is measured in years. Percentile performance scores could then be inferred using the formula:

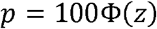

where Φ(·) is the standard Normal cumulative density function. We conducted an independent-samples t-test to compare the percentile performance ranks for the male and female participants who were not vision impaired. There was no significant difference in percentile performance ranks between boys (*M* = 49.5, *SD* = 30.8) and girls (*M* = 48.6, *SD* = 27.7); (*t*(106) = 0.171, *p* = 0.865).

We performed receiver operating characteristic curve analysis on the percentile scores and found the 3.11^th^ percentile was the optimal cut-off for detecting a potential visual search deficit. The area under the curve was 0.957, indicating percentile performance on the TVSAT-C was an excellent parameter for distinguishing between our two groups of participants. We calculated the specificity of the tests to be 97.2%, 95% CI [92.2%, 99.1%] using the 3.11^th^ percentile as the cut-off point. For clarity, the function describing the cut-off point was:

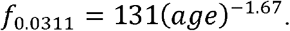

We plotted the visual response time results of the participants with vision impairment (see Methods for individual diagnoses) against their age, and the trajectories of the 5^th^, 50^th^, and 95^th^ percentiles for reference (Figure 3c). All eight vision impaired participants had visual response times that were lower than the average for their age, and seven of the eight participants had visual response times that were lower than the 3.11^th^ percentile. We calculated the sensitivity of the TVSAT-C to be 87.5%, 95% CI [52.9%, 97.8%], under the assumption that each of these participants would have visual search difficulties due to their vision impairment. These results indicated that the TVSAT-C may have good sensitivity for detecting impairment of visual search ability but we require more data from persons with a confirmed visual search deficit to be more confident.

## Discussion

We have developed the TVSAT-C, a real-world test of visual search ability for children that does not require computerised assessment. A normative sample of test results from 108 children between the ages of 3 and 11 demonstrated a smooth developmental trajectory of visual search performance that improved with age following a monomial power law function with an exponent of −1.67. We were able to calculate quantiles, standard z-scores, and define a cut-off score (the 3^rd^ percentile) for typical visual search ability. Using the results from a small sample of vision impaired children with a range of medical conditions, this cut-off gave the TVSAT-C a specificity of 97.2% and sensitivity of 87.5% for discriminating between children with and without vision impairment, indicating it could be a useful tool for detecting visual search deficits in children but further confirmatory studies are required.

The age-dependence of response times observed on the TVSAT-C is likely to result from a combination of factors. Visual search efficiency depends on visual and cognitive abilities such as attentional, perceptual, oculomotor, and executive functions. Additionally, studies have shown that mean response times across a range of tasks decrease throughout childhood and adolescence reflecting improvements in a general processing speed (Kail, 1991; Kail & Ferrer, 2007). Imaging studies have indicated that tract-specific changes in white matter microstructure contribute to improvements in processing speed and executive functions that peak at different ages throughout young adulthood (Peters et al., 2014; Chevalier et al., 2015). Cortical influences on midbrain oculomotor circuitry continue to develop through childhood and adolescence (Johnson, Posner & Rothbart, 1991; Luna, Velanova & Geier, 2008), and eye movement response times decrease from late childhood to adulthood (Luna, Garver, Urban, Lazar & Sweeney, 2004). Other important factors that are likely to contribute to age-dependent changes in test performance include semantic knowledge that has to be learned through domain-specific experience, such as the subtle differences in the visual layout of coins and reading proficiency. However, it is not possible to infer from the current study how much the age-dependent changes in test performance can be explained by changes in specific visual cognitive functions, global development, or learning.

Searching for and identifying letters and coins is an important functional task that children must learn to perform. As such, including the Coins and Letters search tasks in the TVSAT-C was considered worthwhile. However, a potential problem with applying the test battery in younger children is a lack of knowledge and experience of the target letter forms and coin denominations. In the United Kingdom, letter form and coin denomination are formally taught in the first two years of primary school (age: 4-6 years), although this practice varies across countries and cultures. Of course, all goal-oriented visual search tasks require semantic knowledge of key features of the target object to a certain extent (van Gulick, McGugin & Gauthier, 2015), such that the Coins and Letters search tasks are unlikely to assess fundamentally different aspects of visual search ability than the other tasks. Incorporating a range of task difficulties is also useful when testing across a range of ages and levels of education. Whilst younger children should be able to complete most of the simple tasks but struggle on the more difficult tasks, the more difficult tasks may confer greater sensitivity to differences in the ability of older children. The normative data presented here confirm that children younger than 5 years old are often unable to complete the Coins task, and children younger than 4 years old are often unable to complete the Letters task. As such, inability to complete these tasks should not be considered indicative of atypical development or functioning.

Eye tracking has been used previously to better understand visual processing deficits in very young and non-verbal children (Kooiker, Pel, van der Steen-Kant & van der Steen, 2016) and could be used in the future to define the contributions of cognitive functions to impaired performance on the TVSAT-C. Simple eye movement metrics such as the time taken to find and fixate on a target, fixation stability, and the rate of fixation on distractors can be used to infer difficulty with cognitive functions such as shifting attention or suppressing the influence of distractors. More complex metrics involving sequences of eye movements have been related to deficits in visual working memory for previously searched items and locations, or the strategic control of search (e.g. Gilchrist & Harvey, 2000; Husain et al. 2001). As such, eye tracking during functional vision assessment with the TVSAT-C may also be useful for targeting interventions to improve functional vision. For example, abnormal scanning strategies may indicate visual search training as an intervention whereas neglecting to scan to one half of the visual field may indicate hypermetric saccade training instead (during which participants learn to regularly make large angle saccades into the affected hemifield).

The TVSAT-C was designed to assess visual search performance in a realistic setting and does not rely on a computerised display. We believe this gives the TVSAT-C greater ecological validity than standard computerised tests. For example, the field of view over which objects were displayed and had to be searched for in the table-top tasks (we estimate 60 degrees of visual arc) was much wider than that typically occupied by a computerised display (typically 30 degrees at standard viewing distance).

Potential disadvantages of using table-top tests with real objects rather than a digital display is that more time is required to prepare and conduct the TVSAT-C and there is less accuracy as well as potential bias in recording response times. Also, the presence of comorbid motor disorders such as cerebral palsy could make it difficult to interact with physical test materials, reducing performance on the TVSAT-C. We have included objects of a size and shape that are relatively simple to pick up and manipulate in an attempt to counter this, but some items such as small coins may still prove difficult to grasp. The presence of neurological and physical conditions that impair reaching and grasping would need to be considered when interpreting impaired performance on the test. In research, for the purposes of assessing the change in an outcome measure before and after an experimental condition (as opposed to measuring an absolute value), this is less of a concern. In practice, for the purpose of assessing visual ability only, it may be useful to compare performance on the TVSAT-C with a widely used test of visually guided reach and dexterity such as the Box and Blocks Test (Mathiowetz, Federman & Wiemer, 1985; Jongbloed-Pereboom, Nijhuis-van der Sanden & Steenbergen, 2013). Substantially worse performance on the TVSAT-C than the manual dexterity test would indicate that impaired reach and dexterity could not explain the reduced functional vision. Future research including participants with conditions that affect manual dexterity should consider whether other response modes (e.g. pointing, tapping, etc.) could be more appropriate.

The sensitivity of the TVSAT-C requires further evaluation with a larger cohort of vision impaired children who have a confirmed visual search deficit. Both the reliability and validity of the TVSAT-C also require further evaluation. Inter-tester reliability is likely to be good. While there might be some variability in how quickly testers respond on the stopwatch this will be small in comparison to the total time spent searching for the target objects. Test-retest reliability is a potential source for concern. Response times to visual targets are known to be variable from one task to the next. A single response is not a reliable measure, and this is why we included many target objects to search for across the tasks. The external validity of the TVSAT-C could be evaluated simply by comparing results from this test with the results of a standard computerised visual search test from the same participants. It would also be possible to create a computerised version of the TVSAT-C to be displayed on a large, horizontally mounted touch screen device to do so.

The TVSAT-C is a real-world objective measure of visual search ability. The materials required to perform the test are easy and relatively inexpensive to obtain, and the test is simple to run. We would recommend the TVSAT-C as an assessment tool in combination with further investigations. When a vision impairment that could affect visual search ability has been diagnosed or is suspected, the TVSAT-C could be used to infer whether a child has spontaneously learned to compensate for their vision impairment or whether they may require training to improve their visual search ability. We have recently reported on a study in which we used performance on the TVSAT-C to measure improvements in the visual search performance of children and young people with partial visual field loss after they participated in gamified visual search training (Waddington et al., 2018), demonstrating its usefulness as a research tool. Further work to assess the reliability and validity of the test as well as eye tracking investigations of visual search strategies in children will help further develop the TVSAT-C as a potentially useful assessment battery for children with a range of visual and cognitive impairments.

## Acknowledgements

We would like to thank the Summer Scientist Week team for organising the community event at the University of Lincoln where we recruited participants, and Sarah Flynn for her assistance with data collection. We would also like to thank the participating children and families. This work was supported by funding from the WESC Foundation, the Technology Strategy Board (now Innovate UK) and the Medical Research Council of the United Kingdom, as part of a Knowledge Transfer Partnership (Ref: KTP008989). The authors report no conflicts of interest.

## References

Aimola, L., Lane, A., Smith, D., Kerkhoff, G., Ford, G., and Schenk, T. (2014). Efficacy and feasibility of home-based training for individuals with homonymous visual field defects. Neurorehabilitation and neural repair, 28(3), 207–218.

Bartzokis, G., Beckson, M., Lu, P., Nuechterlein, K., Edwards, N., and Mintz, J. (2001). Age-related changes in frontal and temporal lobe volumes in men: a magnetic resonance imaging study. Archives of General Psychiatry, 58(5), 461–465.

Brown, T. (2016). Validity and reliability of the developmental test of visual perception – third edition (DTVP-3). Occupational Therapy in Health Care, 30(3), 272–287.

Brown, T., and Peres, L. (2018a). A critical review of the motor-free visual perception test – fourth edition (MVPT-4). Journal of Occupational Therapy, Schools, & Early Intervention, 11(2), 229–244.

Brown, T., and Peres, L. (2018b). An overview and critique of the test of visual perception skills – fourth edition (TVPS-4). Honk Kong Journal of Occupational Therapy, 31(2), 59–68.

Chesham, A., Gerber, S., Schütz, N., Saner, H., Gutbrod, K., Müri, R., Nef, T., and Urwyler, P. (2019). Search and match task: development of a taskified match-3 puzzle game to assess and practice visual search. JMIR Serious Games, 7(2), e13620. doi: 10.2196/13620.

Chevalier, N., Kurth, S., Doucette, M., Wiseheart, M., Deoni, S., Dean, D., O’Muircheartaigh, J., Blackwell, K., Munakata, Y., and LeBourgeois, M. (2015). Myelination is associated with processing speed in early childhood: preliminary insights. PLoS One, 10(10), e0139897. doi: 10.1371/journal.pone.0139897.

Cooke, D., McKenna, K., Fleming, J., Darnell, R. (2006). Criterion validity of the occupational therapy adult perceptual screening test (OT-APST). Scandinavian Journal of Occupational Therapy, 13(1), 38–48.

Donnelly, S., Hextell, D., Matthey, S. (1998). The Rivermead perceptual assessment battery: It’s relationship to selected functional activities. British Journal of Occupational Therapy, 61(1), 27–32.

Gilchrist, I., and Harvey, M. (2000). Refixation frequency and memory mechanisms in visual search. Current Biology, 10(19), 1209–1212.

Huber, B., Tarasuik, J., Antoniou, M., Garrett, C., Bowe, S., Kaufman, J., and Swinburne Babylab Team. (2016). Young children’s transfer of learning from a touchscreen device. Computers in Human Behavior, 56, 56–64.

Husain, M., Mannan, S., Hodgson, T., Wojcuilik, E., Driver, J. and Kennard, C. (2001). Impaired spatial working memory across saccades contributes to abnormal search in parietal neglect. Brain, 124(5), 941–952.

Huurneman, B., Cox, R., Vlaskamp, B., and Boonstra, F. (2014). Crowded visual search in children with normal vision and children with visual impairment. Vision Research, 96, 65–74.

Ivanov, I., Kuester, S., MacKeben, M., Krumm, A., Haaga, M., Staudt, M., Cordey, A., Gehrlich, C., Martus, P., and Trauzettel-Klosinski, S. (2018). Effects of visual search training in children with hemianopia. PLoS One, 13(7), e0197285. doi: 10.1371/journal.pone.0197285.

Johnson, M., Posner, M., and Rothbart, M. (1991). Components of visual orienting in early infancy: contingency learning, anticipatory looking, and disengaging. Journal of Cognitive Neuroscience, 3(4), 335–344.

Johnson, S. (2019) Development of visual-spatial attention. In Current Topics in Behavioral Neurosciences. Berlin, Heidelberg: Springer. doi: 10.1007/7854_2019_96.

Jongbloed-Pereboom, M., Nijhuis-van der Sanden, M., and Steenbergen, B. (2013). Norm scores of the box and block test for children ages 3-10 years. American Journal of Occupational Therapy, 67(3), 312–318.

Kail, R. (1991). Developmental change in speed of processing during childhood and adolescence. Psychological Bulletin, 109(3), 490–501.

Kail, R., and Ferrer, E. (2007). Processing speed in childhood and adolescence: longitudinal models for examining developmental change. Child Development, 78(6), 1760–1770.

Katz, N., Itzkovich, M., Averbuch, S., and Elazar, B. (1989). Loewenstein occupational therapy cognitive assessment (LOTCA) battery for brain injured patients: reliability and validity. American Journal of Occupational Therapy, 43(3), 184–192.

Klahr, D. (2012). Beyond Piaget: a perspective from studies of children’s problem solving abilities. In A. Slater and P. Quinn (Eds.), Developmental Psychology: Revisiting the Classic Studies (pp. 56–70). London, UK: Sage Publications.

Koenraads Y., van der Linden, D., van Schooneveld M., Imhof S., Gosselaar P., Porro G., and Braun, K. (2014). Visual function and compensatory mechanisms for hemianopia after hemispherectomy in children. Epilepsia, 55(6), 909–917.

Kooiker, M., Pel, J., van der Steen-Kant, S., and van der Steen, J. (2016). A method to quantify visual information processing in children using eye tracking. Journal of Visualized Experiments, 113, 54031. doi: 10.3791/54031.

Kran, B., Lawrence L., Mayer, D., and Heidary, G. (2019). Cerebral/cortical visual impairment: a need to reassess current definitions of visual impairment and blindness. Seminars in Pediatric Neurology, 31, 25–29.

Luna, B., Garver, K., Urban, T., Lazar, N., and Sweeney, J. (2004). Maturation of cognitive processes from late childhood to adulthood. Child Development, 75(5), 1357–1372.

Luna, B., Velanova, K., and Geier, C. (2008). Development of eye-movement control. Brain and Cognition, 68(3), 293–308.

Mathiowetz, V., Federman, S., and Wiemer, D. (1985). Box and Block Test of manual dexterity: norms for 6-19 year olds. Canadian Journal of Occupational Therapy, 52(5), 241–245.

Newcombe, R. (1998). Interval estimation for the difference between independent proportions: comparison of eleven methods. Statistics in Medicine, 17(8), 873–890.

Ong, Y., Jacquin-Courtois, S., Gorgoraptis, N., Bays, P., Husain, M., and Leff, A. (2015). Eye-Search: a web-based therapy that improves visual search in hemianopia. Annals of Clinical and Translational Neurology, 2(1), 74–78.

Pambakian, A., Mannan, S., Hodgson, T., and Kennard, C. (2004). Saccadic visual search training: a treatment for patients with homonymous hemianopia. J Neurol Neurosurg Psychiatry, 75(10), 1443–1448.

Peters, B., Ikuta, T., DeRosse, P., John, M., Burdick, K., Gruner, P., Prendergast, D., Szeszko, P., and Malhotra, A. (2014). Age-related differences in white matter tract microstructure are associated with cognitive performance from childhood to adulthood. Biological Psychiatry, 75(3), 248–256.

Plude, D., Enns, J., and Brodeur, D. (1994). The development of selective attention: A lifespan overview. Acta Psychologica, 86(2-3), 227–272.

Reitan, R., and Wolfson, D. (2004). The trail making test as an initial screening procedure for neuropsychological impairment in older children. Archives of Clinical Neuropsychology, 19, 281–288.

Roth, T., Sokolov, A., Messias, A., Roth, P., Weller, M., and Trauzettel-Klosinski, S. (2009). Comparing explorative saccade and flicker training in hemianopia: a randomized controlled study. Neurology, 72(4): 324–331.

Sahraie, A., Smania, N., and Zihl, J. (2016). Use of NeuroEyeCoach™ to improve eye movement efficacy in patients with homonymous visual field loss. BioMed Research International, 2016, 5186461. doi: 10.1155/2016/5186461.

Schuett, S., Heywood, C., Kentridge, R., Dauner, R, and Zihl, J. (2012). Rehabilitation of reading and visual exploration in visual field disorders: transfer or specificity? Brain, 135(3), 912–921.

Trick, L., and Enns, J. (1998). Lifespan changes in attention: the visual search task. Cognitive Development, 13(3), 369–386.

Turton, A., Angilley, J., Chapman, M., Daniel, A., Longley, V., Clatworthy, P., and Gilchrist, I. (2015). Visual search training in occupational therapy – an example of expert practice in community-based stroke rehabilitation. British Journal of Occupational Therapy, 78(11), 674–687.

van Gulick, A., McGugin, R., Gauthier, I. (2016). Measuring nonvisual knowledge about object categories: The semantic Vanderbilt expertise test. Behavior Research Methods, 48(3), 1178–1196.

van Vugt, P., Fransen, I., Creten, W., and Paquier, P. (2000). Line bisection performances of 650 normal children. Neuropsychologia, 38(6), 886–895.

Waddington, J., Linehan, C., Gerling, K., Hicks, K., and Hodgson, T. (2015, April). Participatory design of therapeutic video games for young people with neurological vision impairment. In CHI ‘15 Proceedings of the 33^rd^ Annual ACM Conference on Human Factors in Computing Systems. Paper presented at CHI 2015: The ACM CHI Conference on Human Factors in Computing Systems, Seoul Republic of Korea (pp.3533–3542). New York, NY: ACM Press.

Waddington, J., and Hodgson, T. (2017). Review of rehabilitation and habilitation strategies for children and young people with homonymous visual field loss caused by cerebral vision impairment. British Journal of Visual Impairment, 35(3), 197–210.

Waddington, J., Linehan, C., Gerling, K., Williams, C., Robson, L., Ellis, R., and Hodgson, T. (2018). Evaluation of Eyelander, a video game designed to engage children and young people with homonymous visual field loss in compensatory training. Journal of Visual Impairment and Blindness, 112(6), 717–730.

Woods, A., Göksun, T., Chatterjee, A., Zelonis, S., Mehta, A., and Smith, S. (2013). The development of organized visual search. Acta Psychologica, 143(2), 191–199.

Zack, E., and Barr, R. (2016). Interactional quality in learning from touch screens during infancy: context matters. Frontiers in Psychology, 7, 1264. doi: 10.3389/fpsyg.2016.01264.

Zihl, J. (1995). Visual scanning behavior in patients with homonymous hemianopia. Neuropsychologia, 33, 287–303.

Zihl, J. (1999). Oculomotor scanning performance in subjects with homonymous visual field disorders. Visual Impairment Research, 1, 23–31.

Zihl, J. (2000). Rehabilitation of visual disorders after brain injury. Hove, UK: Psychology Press Ltd.

